# Cohesin erases genomic-proximity biases to drive stochastic Protocadherin expression for proper neural wiring

**DOI:** 10.1101/2022.03.09.483674

**Authors:** Lea Kiefer, Gabrielle Isabelle F. Servito, Sandy M. Rajkumar, Jennifer Langen, Anna Chiosso, Alexander Buckley, Elizabeth S. Cha, Adan Horta, Michael H. Mui, Daniele Canzio

**Author notes:** These authors contributed equally to this work (Lea Kiefer, Gabrielle Isabelle F. Servito, Sandy M. Rajkumar, Jennifer Langen, Anna Chiosso).

## Abstract

Clustered Protocadherin (Pcdh) proteins act as cell-surface recognition barcodes for neural circuit formation. Neurites expressing the same barcode repel each other, but this mechanism is deployed in two different ways. For instance, convergence of olfactory sensory neuron (OSN) projections requires stochastic expression of distinct Pcdh isoforms in individual cells, while tiling of neural arbors of serotonergic neurons (5-HTs) requires expression of the same isoform, Pcdhαc2. Despite their essential role, however, the molecular mechanisms of cell-type specific Pcdh barcoding remain a mystery. Here, we uncover a new role of cohesin: that of regulating distance-independent enhancer-promoter interactions to enable random Pcdh isoform choice via DNA loop extrusion in OSNs. Remarkably, this step mediates DNA demethylation of Pcdh promoters and their CTCF binding sites, thus directing CTCF to the chosen promoter. In contrast, the uniform pattern of Pcdh expression in 5-HTs is achieved through conventional cohesin-independent, distance-dependent enhancer/promoter interactions, that favor choice of the nearest isoform. Thus, cell-type specific cohesin deployment converts a distance-dependent and deterministic regulatory logic into a distance-independent and stochastic one. We propose that this mechanism provides an elegant strategy to achieve distinct patterns of Pcdh expression that generate wiring instructions to meet the connectivity requirements of different neural classes.

## MAIN TEXT

The establishment of functional neural circuits requires neurons to project to specific brain regions and make appropriate contacts with numerous synaptic partners. Central to this process is the ability of sister neurites (axons and dendrites from the same cell) to recognize and avoid self, a process known as self-avoidance (or isoneural avoidance)^1,2^. Self-avoidance allows branches from the same cell to minimize their overlap while maximizing their territory occupancy^1^. While projecting into a target space, neurons also require instructions to either minimize overlap with their neighboring cells or maximize their coexistence by coalescing with other neurons^3^. These two types of neuronal patterning have been referred to as “tiling” and “convergence”, respectively^1,3^.

To self-avoid, neurons must display unique combinations of cell-surface recognition molecules that function as barcodes, providing a unique identity to individual cells^1^. This molecular strategy of self-avoidance is shared by invertebrates and vertebrates^2^. In vertebrates, the cell-surface barcodes required for self-avoidance are the clustered Protocadherins (Pcdh), a class of transmembrane proteins that engage in highly specific homophilic interactions^4-7^. In mice, there are a total of 116 clustered Pcdh genes, 58 on each of the two parental chromosomes, organized into three tandemly-arranged gene clusters (α, β and γ), which span nearly 1 million base-pairs of genomic DNA^8,9^ (Fig. 1a). Distinct patterns of Pcdh gene expression have been observed in different neural cell types. These patterns have been proposed to carry specific information required by neurons to both regulate their self-avoidance mechanisms and to promote their appropriate connections with neighboring cells^4^. This neuron type-dependent pattern of Pcdh expression is illustrated by recent studies performed in serotonergic neurons (5-HTs) and olfactory sensory neurons (OSNs) in mice (Fig. 1a). Specifically, all 5-HTs primarily express a single Pcdh gene (Pcdhαc2), therefore displaying an identical Pcdh barcode on their cell-surfaces (Fig. 1a)^10,11^. This pattern of Pcdh expression not only causes neurites from individual 5-HTs to self-avoid but also promotes repulsion between neighboring 5-HTs^10,11^. This overall behavior allows 5-HTs to “tile” as they project from the dorsal raphe to distal regions of the brain^10,11^. Deletion of Pcdhαc2 results in a striking tiling defect of 5-HT projections^10,11^, a self-avoidance phenotype similar to that previously observed in starburst amacrine cells upon genetic removal of the Pcdhγ genes^12^. In 5-HTs, this wiring defect results in a non-uniform release of serotonin, which is likely responsible for the behavioral abnormalities observed in Pcdhα gene cluster knock-out mice^10,11^. Contrary to 5-HTs, olfactory sensory neurons provide an example of an orthogonal Pcdh expression profile as individual OSNs do not express a single Pcdh isoform but rather a repertoire of Pcdh genes (Fig. 1a)^13^. This repertoire appears to be generated by a complex mechanism of stochastic Pcdh promoter choice and ensures that essentially every OSN displays a distinct barcode on its cell surface^13^. This vast Pcdh cell surface diversity is required for the convergence of OSN axons expressing the same olfactory receptor (OR) into individual glomeruli of the olfactory bulb (OB)^13-16^. Overriding Pcdh diversity in OSNs by expressing high levels of specific Pcdh isoforms leads to axon convergence defects and disruption of glomeruli formation in the OB^13^. Consequently, these mice are unable to effectively discriminate between odors^13^.

**Figure 1.**
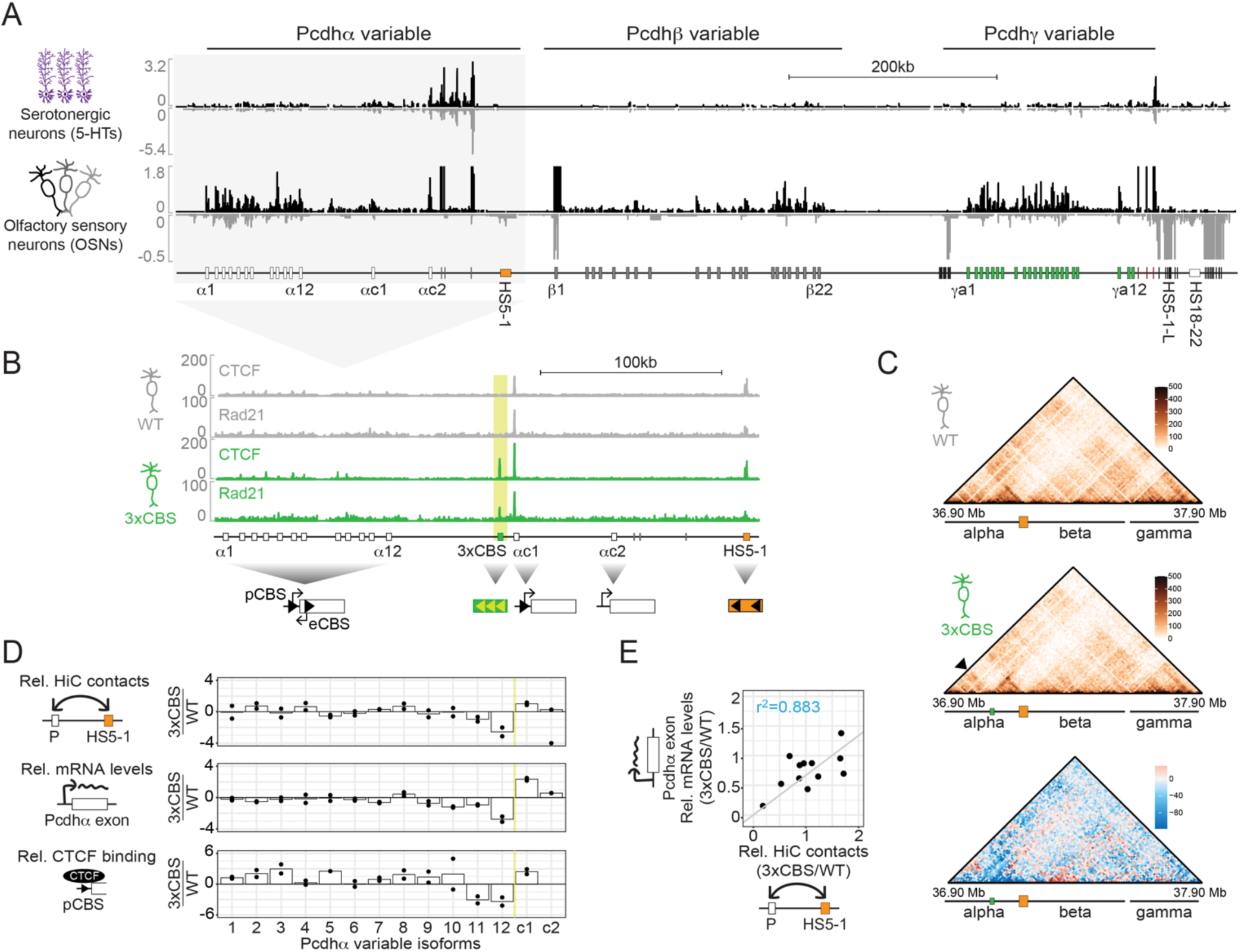
Cohesin extrusion through the Pcdhα gene cluster randomizes promoter choice. **(a)** Genomic architecture of clustered Pcdh genes and RNA expression profiles of 5-HTs^41^ and OSNs **(b)** ChIP-seq profiles for CTCF and Rad21 from total olfactory epithelium from WT and 3xCBS mice. Ectopic 3xCBSs are indicated in yellow, the orientations of the CBSs in bold arrows. **(c)** HiC contact maps at 10kb resolution for the Pcdh gene cluster in WT and 3xCBS OSNs and their difference contact map. Arrowhead indicates the newly formed boundary. **(d)** Fold change between 3xCBS and WT OSNs of HiC contacts between the HS5-1 enhancer and the variable Pcdhα genes (Top), RNA expression for the Pcdhα variable exons (Middle), and CTCF occupancy quantified from ChIP-seq (Bottom). Yellow line indicates the location of the 3xCBS. **(e)** Correlation plot of the RNA expression of Pcdhα alternate exons and HiC contact between the alternate promoters and the HS5-1 enhancer in 3xCBS relative to WT OSNs.

These studies suggest that the generation of distinct Pcdh expression patterns in different neural cell types is critical to the establishment of complex cell-type specific neural circuits and animal behavior. Yet, how cell-type specific Pcdh expression patterns are generated from the same gene cluster remains a mystery.

### Cohesin extrusion in the Pcdhα gene cluster randomizes promoter choice

The mouse Pcdhα gene cluster bears 14 “variable” 5’ exons, each driven by its own promoter (Fig. 1a). Pcdhα 1-12 are also referred to as “alternate”, while Pcdhαc1 and Pcdhαc2 are referred to as “c-type”. Downstream of the c-type exons, there are three small “constant” exons encoding most of the intracellular domain shared between all Pcdhα isoforms (Fig. 1a)^17^. Previous studies have shown that stochastic Pcdhα promoter choice in OSNs requires transcription of an antisense long non-coding RNA (as-lncRNA) initiated from a promoter located within each Pcdhα 1-12 coding region^18^. Transcription of the as-lncRNA through the sense promoter leads to DNA demethylation of the chosen promoter and its CTCF binding sites (CBSs)^18^. This step is coupled to CTCF binding to its CBSs, long-range DNA interactions between the chosen Pcdhα promoter and a distant transcriptional enhancer, HS5-1, by cohesin and transcriptional activation of the chosen promoter (Fig. 1b)^18-22^.

Enhancer/promoter interactions mediated by CTCF and cohesin have been proposed to result from actively extruding cohesin molecules on DNA which are preferentially stalled by CTCF at convergently oriented CBSs^23-25^. In the context of the Pcdhα gene cluster, Pcdhα 1-12 promoters/exons bear two CBSs (pCBS and eCBS, respectively), while Pcdhαc1 solely has a pCBS (Fig. 1b). In contrast, Pcdhαc2 lacks a CBS (Fig. 1b). As in the case of Pcdhα 1-12, the HS5-1 enhancer bears two CBSs, whose orientations are convergent to those of Pcdhα 1-12 (Fig. 1b). To test if stochastic Pcdhα promoter choice in primary neurons is mediated by DNA loop extrusion, we engineered a mouse bearing an insertion of three ectopic CTCF binding sites (3xCBS) upstream of the Pcdhαc1 promoter in the same orientation as the sites in the HS5-1 enhancer (Fig. 1b and Extended Data Fig. 1a). ChIP-Seq analysis demonstrated enrichment of both CTCF and cohesin to these sites from cells isolated from the mouse olfactory epithelium (OE) (Fig. 1b; the cohesin subunit, Rad21, was used to assay cohesin localization). Interestingly, the ChIP-Seq data also revealed a decrease in stalling of cohesin at the HS5-1 enhancer (Fig. 1b). Both observations suggest that the three CBSs stall cohesin extrusion through the gene cluster. To investigate this possibility, we performed in situ Hi-C experiments using OSNs (obtained by FACS, see methods for details) bearing the 3xCBS insertion. Consistent with our ChIP-Seq data, a new domain boundary was formed upstream of the Pcdhαc1 gene (Fig. 1c). Quantification of the Hi-C data revealed a distance-biased loss of enhancer/promoter contacts, whereby the promoters most proximal to and upstream of the newly formed boundary were most affected (Fig. 1d, Top). To test the functional consequence of this newly formed boundary on Pcdh gene expression, we performed RNA-Seq in OSNs. Strikingly, we observed a similar distance bias in Pcdhα 1-12 expression in the presence of the 3xCBS sites (Fig. 1d, Middle). Transcription from the Pcdhαc1 promoter instead increased (Fig. 1d and Extended Data Fig. 1b). Thus, there is a clear correlation between transcription and promoter/enhancer interaction (Fig. 1e).

Finally, given the previously established strong correlation between transcription of Pcdh promoters, their contacts with the HS5-1 enhancer and their occupancy by CTCF and cohesin^18-22^, we quantified the binding of CTCF to the pCBSs of each variable exon in WT and 3xCBS OSNs. Remarkably, consistent with loss of RNA expression and enhancer/promoter contacts, binding of CTCF to the pCBS of Pcdhα 11 and 12 was severely reduced (Fig. 1d, Bottom).

### Cohesin extrusion promotes DNA demethylation of Pcdhα promoters

We next wondered whether the observed loss of CTCF binding at the Pcdhα12 promoter was due to DNA hypermethylation of its pCBS. To test this, we employed whole-genome bisulfite sequencing (WGBS) studies and observed an increase in 5-methylcytosine (5mC) at both the promoter and the CpG element at position 12 of the canonical CTCF binding motif in the pCBS of Pcdhα12 (Fig. 2a and Extended Data Fig. 1c), a modification known to impact CTCF binding to DNA^26^.

**Figure 2.**
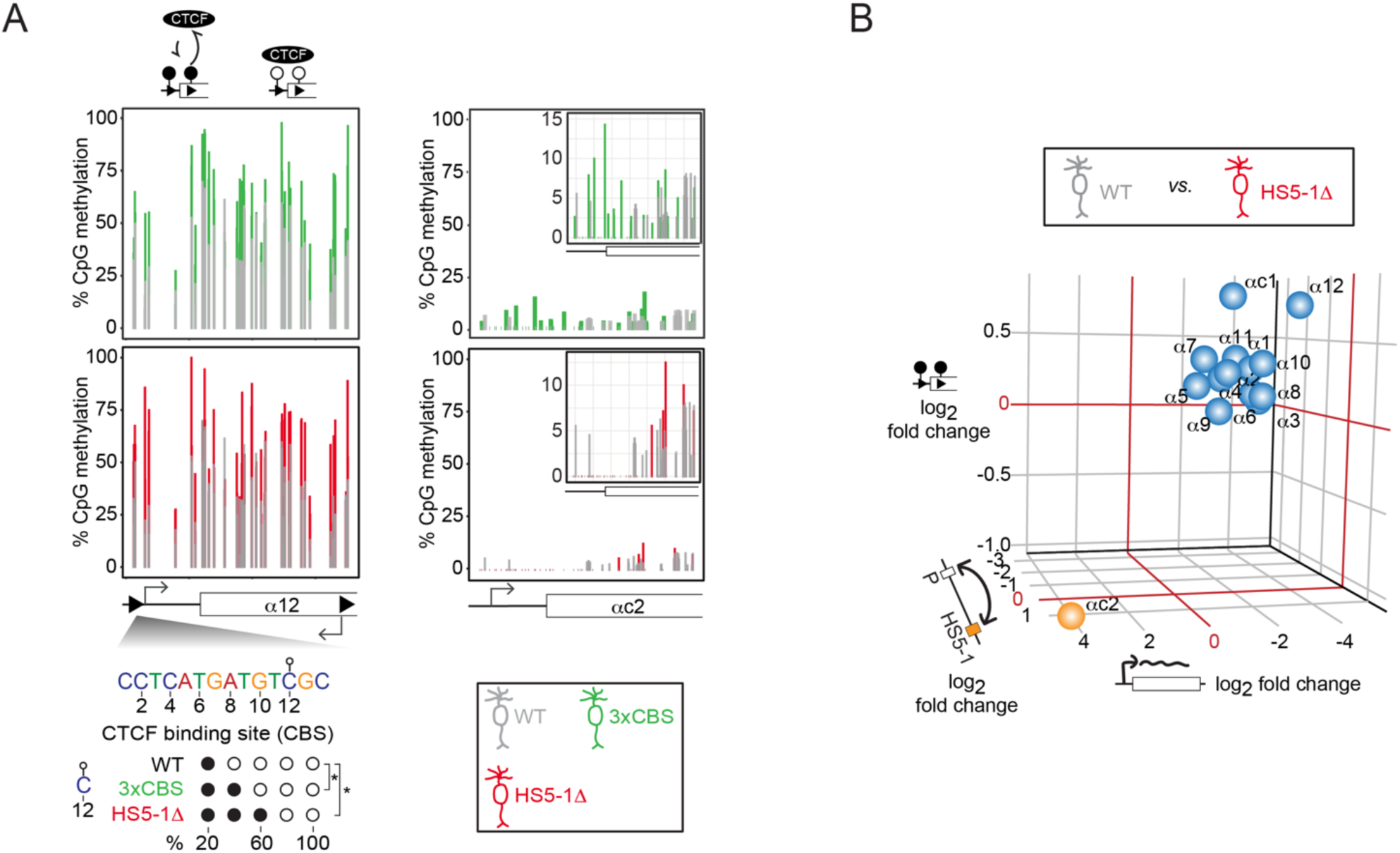
Enhancer/Promoter contacts induce DNA demethylation of Pcdhα promoters. (**A**) **(a)** Percent CpG methylation across the Pcdhα12 promoter in WT (grey), 3xCBS (green) and HS5-1Δ (red). Two biological replicates shown as bars per individual cytidine in the indicated Pcdhα promoters with the methylation status of cytosine 12 of the Pcdhα12 promoter CTCF binding motif in WT, 3xCBS and HS5-1Δ OSNs (WT 19%, 3xCBS 42%, HS5-1Δ 63%). Inserts show zoomed in methylation levels for Pcdhαc2 promoter. **(b)** Log2 fold-change of Pcdhα isoform expression by RNA-seq (x-axis), interaction with the HS5-1 enhancer by in situ HiC (z-axis) and promoter methylation levels by WGBS (y-axis) in HS5-1Δ compared to WT OSNs.

Altogether, these data suggest that DNA loop extrusion by cohesin regulates Pcdhα promoter choice by mediating enhancer/promoter contacts to allows for DNA demethylation of Pcdh promoters. To test this possibility, we engineered a mouse line bearing a deletion of the HS5-1 enhancer and its CBSs (HS5-1Δ) (Extended Data Fig. 2a) and performed WGBS in HS5-1Δ OSNs. Consistent with our hypothesis, deletion of the HS5-1 enhancer increased the level of 5mC for all Pcdhα promoters relative to WT OSNs (Extended Data Fig. 2b). The sole exception was Pcdhαc2, whose ground state is hypomethylated, and the low level of 5mC was further reduced upon deletion of the HS5-1 enhancer (Fig. 2a and Extended Data Fig. 2b). Importantly, consistent with the role of DNA loop extrusion in promoting Pcdh promoter demethylation, we observed an even higher increase in 5mC levels of position 12 of the pCBS of Pcdhα12 (Fig. 2a).

To investigate the interplay between long-range HS5-1 enhancer/promoter contacts, Pcdhα transcription and promoter CpG methylation, we performed in situ Hi-C and RNA-seq studies in HS5-1Δ and WT OSNs. Consistent with the increase in promoter DNA methylation, deletion of the HS5-1 sequence resulted in a severe decrease of long-range contacts between the 20kb region surrounding the HS5-1 enhancer and the Pcdhα 1-12 and Pcdhαc1 promoters (Fig. 2b). Similarly, we observed a near-complete loss of both sense and antisense RNA transcription of Pcdhα 1-12 and Pcdhαc1 (Fig. 2b and Extended Data Fig. 2c,d). Strikingly, however, the Pcdhαc2 gene behaved differently from the rest of the cluster: RNA expression increased by about 20-fold (Fig. 2b and Extended Data Fig. 2d) and its promoter gained contacts with the DNA sequence nearby the HS5-1 enhancer (Fig. 2b and Extended Data Fig. 2d). Finally, deletion of the HS5-1 enhancer also resulted in an increase in transcription of the Pcdhβ genes located most proximal to the Pcdhα gene locus (Extended Data Fig. 2d). No effect was observed on the transcription of Pcdhβ genes located more distant to the Pcdhα cluster or of the Pcdhγ genes (Extended Data Fig. 2d).

Taken together, these data suggest that Pcdhα promoter choice is regulated by DNA loop extrusion and that cohesin activity drives a functional switch between two orthogonal transcriptional programs: one in which Pcdhα promoter choice is stochastic and enhancer distance-independent, and one in which promoter choice is non-random and distance-biased, where the sole chosen promoter is that of Pcdhαc2.

### Pcdhα promoters are regulated by distinct regulatory sequences that have contrasting requirements for DNA loop extrusion

The observation that Pcdhαc2 expression is upregulated upon deletion of HS5-1 suggests the activity of a separate regulatory sequence specific to the promoter of Pcdhαc2. In search of such a sequence, we examined the Pcdhα gene cluster for novel regulatory elements by performing ChIP-Seq in OSNs, focusing on H3K4me3 and H3K27ac indicative of transcriptional activation, and absence of the transcriptional repressive mark H3K9me3. Analysis of our ChIP-Seq datasets along with published ATAC-Seq data in OSNs^27^ revealed several putative regulatory elements, in addition to HS5-1 (Fig. 3a). Of particular interest were two putative regulatory sequences located downstream of the Pcdhαc2 gene and upstream of the HS5-1 enhancer, which we designated as HS8-7 (Fig. 3a). The HS7 sequence was previously identified as a Pcdhα transcriptional enhancer required for the expression of the most enhancer-proximal Pcdhα promoters in different whole brain mouse tissues^22,28^. Our ChIP-Seq data showed that, in contrast to HS5-1, the HS8-7 putative enhancer elements are not bound by either CTCF or cohesin (Fig. 3a). To test the enhancer function of the HS8-7 sequence, we next engineered a mouse in which the HS8-7 sequence is deleted (HS8-7Δ) (Extended Data Fig. 2a). Consistent with its role as a transcriptional enhancer in OSNs, deletion of the HS8-7 sequences resulted in a dramatic loss of expression of Pcdhαc2, Pcdhαc1 and Pcdhα12 (Fig. 3b and Extended Data Fig. 2d). These results differ from the impact of deleting HS5-1, thus suggesting that the two regulatory sequences display distinct promoter specificities. To test this, we measured the extent of their contacts with all Pcdhα promoters and observed that, contrary to the HS5-1 enhancer, the HS8-7 enhancer displays a distance bias towards the closest Pcdhαc2 promoter (Fig. 3c).

**Figure 3.**
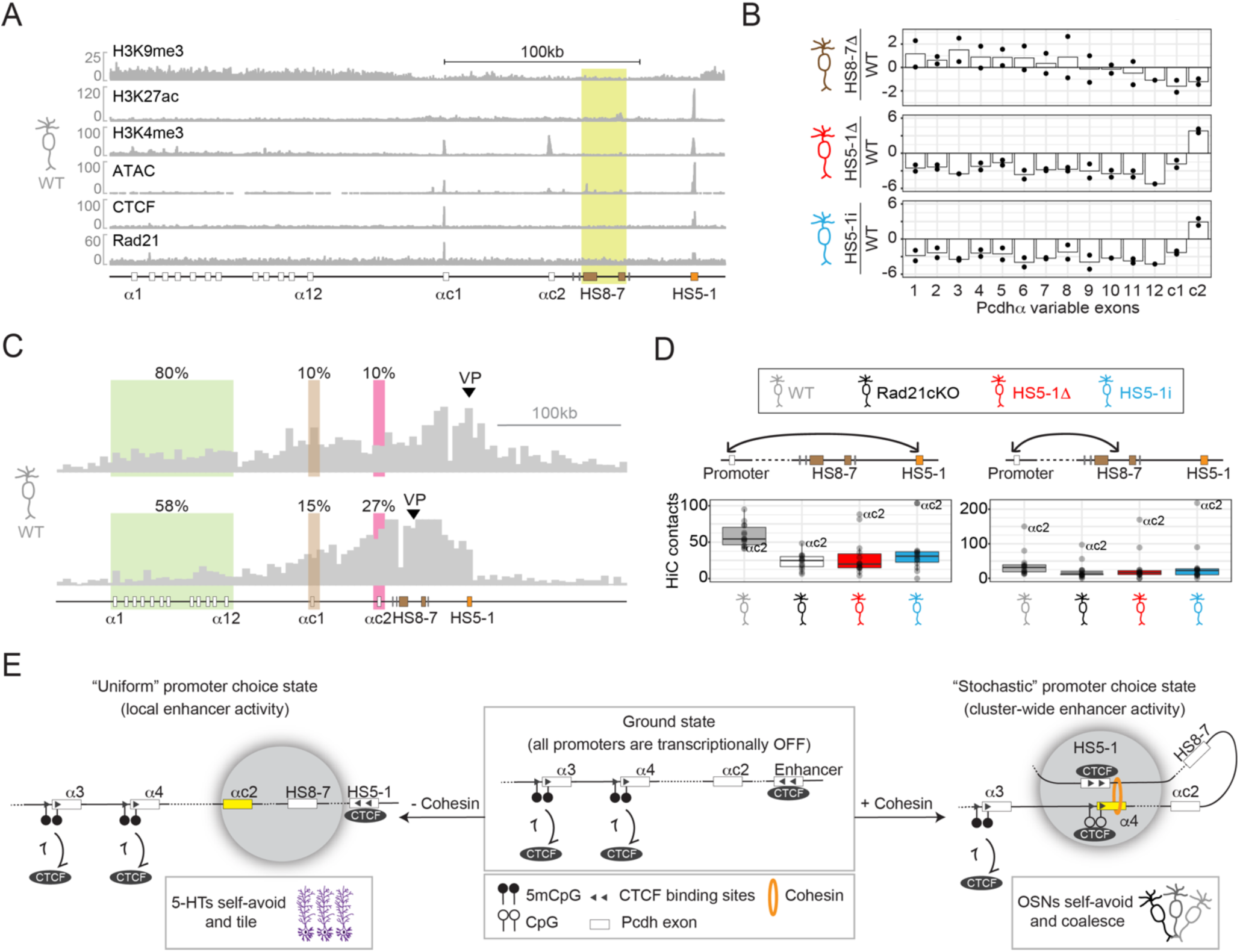
Pcdh promoters are regulated by the activities of distinct regulatory sequences and their dependence on cohesin extrusion. **(a)** Fixed-ChIP-Seq, Native-ChIP-Seq and ATAC-Seq profiles of the Pcdhα cluster in OSNs. **(b)** Fold-change of mRNA levels of Pcdhα isoforms (dots indicate biological replicates). Data from HS8-7Δ, HS5-1Δ and HS5-1i OSNs. **(c)** Virtual 4C of the chromatin contacts from the HS5-1 and the HS8-7 enhancers. Viewpoint indicated by arrowheads. Quantification by percent contacts. **(d)** Quantification of chromatin contacts between the HS5-1 and the HS8-7 enhancer with the Pcdhα promoters in WT, Rad21cKO, HS5-1Δ and HS5-1i OSNs. **(e)** Model: In their ground state, all alternate Pcdhα promoters are methylated, preventing CTCF binding. Expression of the sole Pcdhαc2 isoform is DNA loop extrusion independent and requires the local activity of the HS8-7 enhancer. This mechanism allows axon tiling of 5-HTs. Stochastic Pcdhα promoter activation involves cohesin-mediated contacts with the HS5-1 enhancer which couples demethylation of the chosen promoter to CTCF binding. In OSNs, this mechanism generates sufficient protein isoform diversity required for OSN axons to coalesce.

### DNA loop extrusion counteracts promoter choice biases driven by genomic proximity

Based on the above data, we next wondered whether DNA loop extrusion governs the differential promoter specificities of the HS5-1 and HS8-7 enhancers and engineered a mouse line in which we inverted the HS5-1 enhancer (HS5-1i, Extended Data Fig. 2a). Remarkably, RNA-seq revealed a near-complete loss of Pcdhα 1-12 and Pcdhαc1 expression accompanied by a dramatic upregulation of Pcdhαc2 expression (Fig. 3b and Extended Data Fig. 2d). As in the case of the HS5-1Δ mouse, we also observed loss of Pcdhα as-lncRNA transcription (Extended Data Fig. 2c), and an increase in Pcdhβ expression of the genes located proximal to Pcdhα gene cluster (Extended Data Fig. 2d). Thus, the observation that inversion of the HS5-1 enhancer phenocopies its deletion suggests that its activity is dependent on cohesin-mediated DNA loop extrusion.

To investigate the structural basis of this transcriptional switch, we performed in situ Hi-C experiments in HS5-1i OSNs and compared them to HS5-1Δ and to OSNs where the cohesin subunit Rad21 was conditionally deleted (Rad21cKO) (Extended Data Fig. 3a). Consistent with a role of DNA loop extrusion in the regulation of the HS5-1 enhancer activity, all three perturbations resulted in loss of contacts between the HS5-1 enhancer and the Pcdhα alternate promoters, and a gain of interactions between the HS5-1 and HS8-7 enhancers with the Pcdhαc2 promoter (Fig. 3d and Extended Data Fig. 3b). Interestingly, deletion and inversion of the HS5-1 enhancer also resulted in increased contacts of the region surrounding the HS5-1 enhancer and the Pcdhβ cluster, but not the Pcdhγ cluster (Extended Data Fig. 3b). This effect was not seen in the Rad21cKO case and provides a mechanistic explanation for the increase in Pcdhβ transcription observed by RNA-Seq (Extended Data Fig. 2d).

Taken together, these data suggest that cohesin-mediated DNA loop extrusion is required to distribute the regulatory activity of the distal HS5-1 enhancer throughout the Pcdhα cluster to mediate random escape of Pcdhα alternate promoters from silent chromatin (Fig. 3e, Right). In the absence of DNA loop extrusion, the activity of the HS8-7 enhancer is biased to the nearest Pcdh promoter, Pcdhαc2 (Fig. 3e, Left). Thus, DNA loop extrusion defines the probability of Pcdh promoter choice by turning a process that is, by default, distance-biased to one that is distance-independent (Fig. 3e). According to this model, differential regulation of cohesin activity on DNA results in the generation of distinct levels of Pcdh isoform barcoding diversity, which is the basis of cell-type specific mechanisms of neural arborization. Consistent with this is our observation that ablating DNA loop extrusion by either deletion or inversion of the HS5-1 enhancer reconfigures the Pcdhα transcriptional profile of OSNs to that of 5-HTs, whereby every neuron uniformly expresses only Pcdhαc2. Thus, this model predicts that, in the context of 5-HTs, axon tiling relies solely on the cohesin-independent and local activity of the HS8-7 enhancer driving expression of Pcdhαc2 (Fig. 3e). On the contrary, in the context of OSNs, DNA loop extrusion generates sufficient single neuron Pcdh cell-surface diversity required for proper OSN axon convergence (Fig. 3e). Next, we sought to test the mechanistic and physiological predictions of this model in 5-HTs and OSNs.

### Cohesin-independent enhancer activity drives Pcdhαc2-only expression and axon tiling of serotonergic neurons

To test our model in serotonergic (5-HT) neurons, we first performed RNA-Seq from the raphe isolated from WT and HS8-7Δ mice and observed a loss of Pcdhαc2 expression upon deletion of HS8-7 (Fig. 4a,b). To further probe the promoter specificities of the HS8-7 and HS5-1 enhancers in 5-HTs at the single neuron level, we performed RNA Fluorescence in situ hybridization (FISH) experiments. Remarkably, consistent with our model and with our data from OSNs (Fig. 3b), deletion of the HS8-7 enhancer resulted in a reduction of Pcdhαc2 mRNA levels while deletion of the HS5-1 resulted in upregulation (Fig. 4c and Extended Data Fig. 4a). Next, we asked whether deletion of the HS8-7 enhancer results in 5-HT wiring defects by examining the ability of 5-HT axons to project to the hippocampus, and observed that 5-HT axon terminals failed to properly arborize in HS8-7Δ mice (Fig. 4a,d). Strikingly, this defect phenocopied that observed upon deletion of the Pcdhαc2 gene^10,11^. Importantly, consistent with the model that Pcdhαc2 expression is regulated independently of the chromatin architecture of the cluster imposed by DNA loop extrusion, we did not observe a similar phenotype in HS5-1Δ, HS5-1i, 3xCBS mice (Fig. 4d and Extended Data Fig. 4b).

**Figure 4.**
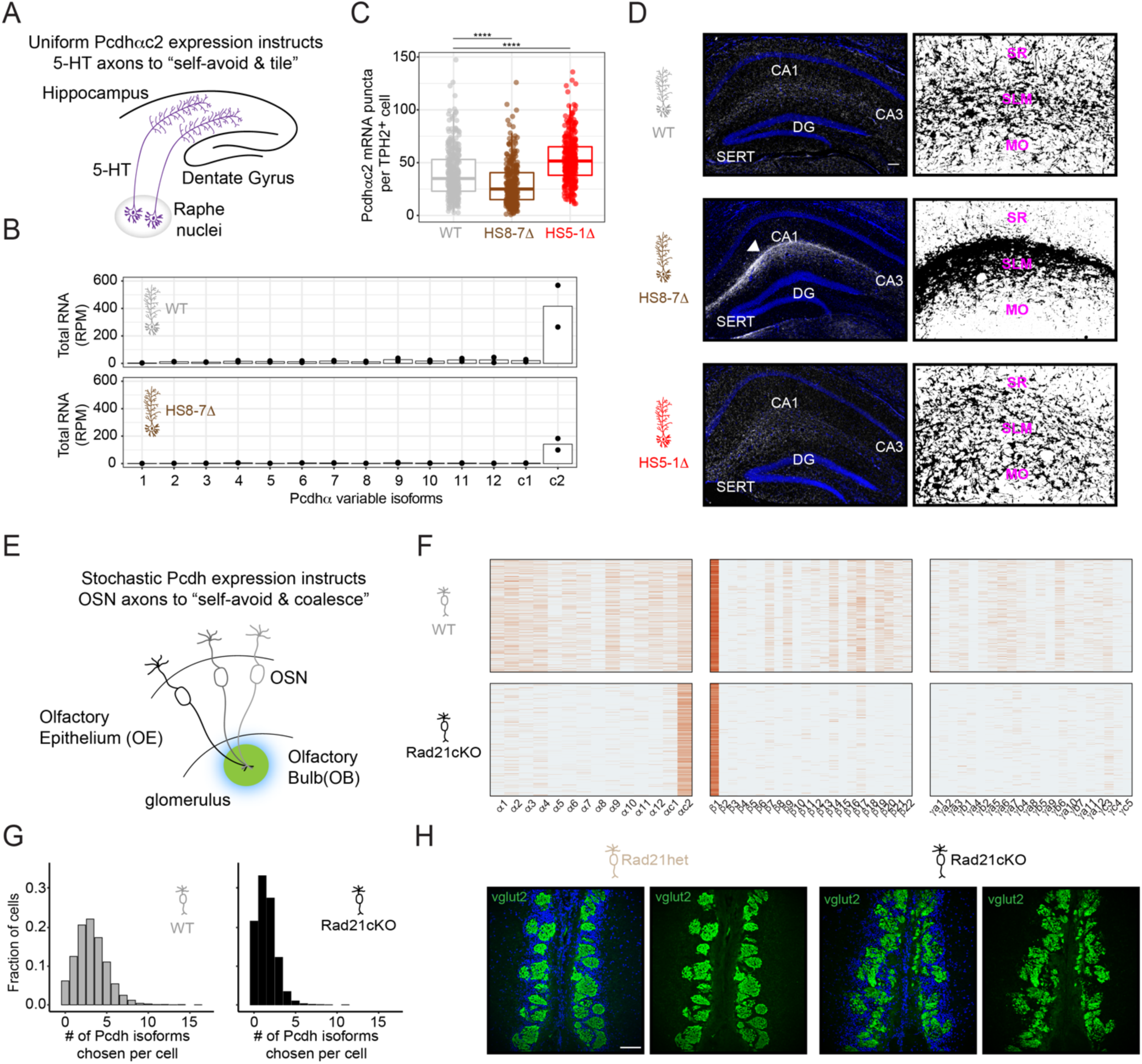
Cohesin-independent and dependent Pcdh expression mechanisms regulate 5-HT tiling and OSN coalescence. **(a)** Schematic of 5-HT axon tiling from the raphe to the hippocampus. **(b)** RNA-seq levels of Pcdhα isoforms detected in the dorsal raphe of HS8-7Δ and WT mice (two biological replicates each). **(c)** Quantification of number of Pcdhαc2 mRNA puncta in TPH2^+^ cells. **(d)** 5-HT wiring defects in HS8-7Δ mice but not in HS5-1Δ relative to WT. Left: 5-HTs are indicated by IHC against SERT. SERT, white; DAPI, blue. Arrow indicates clumping of 5-HT axons. CA1, CA3 and DG refer to cornu ammonis 1 and 3 and dentate gyrus, respectively. Right: Binary mask of SERT signal in presented images. SR, SLM and MO refer to stratum radiatum, stratum lacunosum moleculare, and molecular layer. **(e)** Schematic of OSN axon projection from the OE and coalescence in the OB. **(f)** Single-cell Pcdh promoter choice from all three clusters from individual OSNs isolated from WT and Rad21cKO. Each row indicates a single cell, each column a single isoform. **(g)** Histogram of number of chosen Pcdh promoters per individual cell in WT and Rad21cKO OSNs. **(h)** IHC against Vglut2 in coronal sections in ompcre;Rad21(fl/wt);tdTomato(fl/fl) (Rad21het) and in ompcre;Rad21(fl/fl);tdTomato(fl/fl) (Rad21cKO). Vglut2, green; DAPI, blue. Scale bars: 100 μm.

### Cohesin-dependent enhancer activity generates Pcdh isoform diversity required for proper axon wiring in OSNs

Our model predicts that disruption of DNA loop extrusion in OSNs abolishes Pcdh isoform diversity by switching from stochastic Pcdh promoter choice to uniform choice of Pcdhαc2 at a single cell level and that such effect would have implications for the ability of OSN axon projections to establish appropriate glomeruli structures (Fig. 4e). As bulk RNA-seq can only reveal expression levels of a mixed population of cells, we turned to single cell RNA-seq to query Pcdh gene choice at a single cell level in 3xCBS, HS5-1i and Rad21cKO OSNs. Consistent with our model, we observed a dramatic loss of Pcdhα isoform diversity in OSNs isolated from all three genotypes (Fig. 4f and Extended Data Fig. 4c). Remarkably, impairment of DNA loop extrusion by inversion of the HS5-1 enhancer and removal of cohesin by ablation of Rad21 turned a stochastic promoter choice mechanism into one where the promoter of Pcdhαc2 was chosen most frequently by individual OSNs (Fig. 4f and Extended Data Fig. 4c). Importantly, ablation of Rad21 also led to loss of Pcdhβ and Pcdhγ promoter choice, reducing the overall Pcdh isoform diversity and suggesting that the Pcdhβ and Pcdhγ clusters also require DNA loop extrusion mechanisms for their stochastic expression (Fig. 4f,g). Given the observed loss of Pcdh diversity, we next tested whether ablation of Rad21 would result in any phenotypic consequences. Remarkably, we observed a severe disruption of proper glomeruli structures throughout the entire olfactory bulb in Rad21cKO mice compared to their heterozygous littermates (Rad21het) (Fig. 4h and Extended Data Fig. 4d). While we cannot exclude that this effect is due to additional changes in gene expression upon Rad21 knockout, it is important to note that olfactory receptor gene expression, a key determinant of OSN axon positioning, is not affected (Extended Data Fig. 4e). These data are consistent with the established paradigm that Pcdh protein isoform diversity plays a critical role in the assembly of olfactory circuits^13-16^, and suggest that DNA loop extrusion by cohesin generates the requisite Pcdh barcoding diversity in OSNs by driving stochastic Pcdh promoter choice.

## DISCUSSION

Neural subtype-specific Pcdh expression patterns play a critical role in instructing neural wiring mechanisms. We show that the orthogonal Pcdh expression patterns of olfactory sensory neurons (OSNs) and serotonergic neurons (5-HTs) leverage distinct regulatory elements encoded in the Pcdh gene locus. While uniform Pcdhαc2 protein barcoding in 5-HTs is achieved by the activity of the cohesin-independent proximal HS8-7 enhancer, the generation of Pcdh diversity in OSNs requires the activity of the cohesin-dependent distal HS5-1 enhancer. In the context of OSNs, we propose that deployment of cohesin-mediated DNA loop extrusion serves, at least, two functions: (1) it provides the means of directing the activity of the HS5-1 enhancer to distant promoters, located as far as 200,000 base-pairs away, thus rendering a locus that by default functions as enhancer-distance dependent into one that is enhancer-distance independent, and (2) it equalizes the probability of any of the distant Pcdh promoters to be transcriptionally chosen (Fig. 3e). The latter function is achieved by cohesin coupling long-range HS5-1 enhancer/promoter contacts to DNA demethylation of the promoters and their CBSs (Fig. 3e). Demethylation of the CBSs allows for CTCF binding, which we hypothesize locks in Pcdh promoter choice (Fig. 3e). As the pattern of Pcdh promoter choice in numerous established cell lines appears stable^20,29^, it is possible that Pcdh promoter demethylation is epigenetically maintained for the lifetime of a terminally differentiated neuron.

Supporting our proposed model is the surprising observation that disruption of cohesin activity in OSNs results in a Pcdh transcriptional profile of a different class of neurons – serotonergic neurons. Considering that a human brain houses nearly 100 billion neurons, it is likely that the Pcdh expression profiles of OSNs and 5-HTs represent two examples of a diverse and complex spectrum of Pcdh transcriptional profiles. Indeed, mouse Purkinje neurons express yet a different Pcdh transcriptional profile from the ones characterized here^30,31^. We speculate that neural subtypes have evolved a regulatory logic to achieve distinct patterns of Pcdh gene expression by tuning the DNA loop extrusion activity of cohesin to leverage the complex architecture of the Pcdh gene cluster and its regulatory elements. Interestingly, there is precedence for such a process in B-cells, which tune their cohesin activity to maximize antibody diversity via V(D)J recombination during maturation^32,33^. Considering that Pcdh expression profiles influence neural wiring and function, we propose that this logic provides an elegant strategy to overcome the daunting challenge of generating specific wiring instructions that meet the connectivity requirements of different neural classes.

Finally, as genetic variants of the cohesin complex and its regulators are associated with cohesinopathies, including Cornelia de Lange Syndrome and Roberts Syndrome^34^, it is reasonable to speculate that the severe intellectual impairments associated with these disorders arise, at least in part, from the dysregulation of clustered Pcdh expression. Consistent with this is the recent observation that clustered Pcdh genes are dysregulated in cells derived from Cornelia de Lange Syndrome patients^35^.

## METHODS

### Animals

Mice were treated in compliance with the rules and regulations of IACUC under protocol number AN-170364-03F. Both male and female animals of age between 4 and 16 weeks were used for all experimental procedures. All unpublished lines (HS5-1i, HS5-1Δ, HS8-7Δ and 3xCBS) were generated using CRISPR-mediated knock-out and knock-in. The DNA sequence utilized to engineer the 3xCBS mouse line was previously tested to be functional in cell culture systems ^36^. Primary FACS-sorted cells were obtained from dissected main olfactory epithelium. WT, HS5-1i, HS5-1Δ, HS8-7Δ and 3xCBS OSNs were sorted from OMP-ires-GFP mice (OMP = olfactory mature marker gene). Rad21 conditional knockout OSNs were achieved as described previously ^18^. Briefly, Rad21 conditional allele mice were crossed to OMP-ires-Cre mice (Omp^tm1(cre)Jae^). Recombined cells were purified by including a Cre-inducible tdTomato allele (ROSA26-tdtomato, Gt(ROSA)26Sor^tm14(CAG-tdTomato)Hze/J^) in the cross and selecting tdTomato positive cells by FACS. In the text and the figures, we refer to the Rad21 conditional knockout in OSNs as Rad21cKO for homozygous deletion of the floxed allele and as Rad21het for heterozygous deletion of the floxed allele.

### Fluorescence activated cell sorting of mouse OSNs

Cells were dissociated into a single-cell suspension by incubating freshly dissected main olfactory epithelium with papain for 30-40 minutes at 37°C according to the Worthington Papain Dissociation System. Following dissociation and filtering through a 35 μm cell strainer, cells were resuspended in 1X PBS with 2% FBS with DNase (0.0025% final concentration) and DAPI. For *in situ* Hi-C and ChIP-Seq experiments, upon dissociation, cells were fixed with 1% formaldehyde for 10 minutes at room temperature. Formaldehyde was quenched by adding glycine to a final concentration of 0.125 M for 5 minutes at room temperature. Cells were then washed once and resuspended in cold PBS with 2% FBS and DNase (0.0025% final concentration). Fluorescent cells were then sorted on a BD Aria II.

### RNA isolation and sequencing studies

RNA was isolated from tissue using TRIzol. Cell lysate was extracted with bromo-chloropropane and RNA was precipitated with 100% isopropanol supplemented with 10 μg of glycoblue for 10 min at room temperature and then pelleted at 16,000 x g for 30 min at 4°C. The RNA pellet was washed once with 75% ethanol and then resuspended in RNase-free water to a maximal concentration of 200 ng/μl. Genomic DNA contaminants were removed by Turbo DNase. Removal of Turbo DNase was performed by phenol:chloroform extraction and RNA was precipitated as described above and resuspended in RNase-free water and stored at -80°C.

RNA was isolated from coronal sections of the Raphe (20 μm thick) obtained from paraformaldehyde-fixed brain tissue (described in more detail below, see “Preparation of Mouse Brain Tissue”). Slides were washed three times for 5 minutes each with 1X PBS. After the washes, a portion of each section containing approximately the Raphe region was extracted. Next, the RNA was isolated using RecoverAll™ Total Nucleic Acid Isolation Kit for FFPE (Invitrogen) as instructed by the manufacturer.

Sequencing libraries for total RNA were made using the SMARTer Stranded Total RNA-Seq Pico input mammalian RNA kit v2. The quality and quantity of all the libraries was assessed by bioanalyzer and qubit. Libraries were sequenced on a NEXT-Seq 500/550 (UCSF Gladstone Genomic Core).

### Single-cell RNA sequencing studies

Cells were dissociated from the main olfactory epithelium from 8 to 12 weeks old female mice as described above. About 10,000 cells were FACS purified for OSNs and submitted for 10x Genomics GEM generation using the Single Cell 5’ Gene Expression set up and sequenced using a NovaSeq SP100 PE50 (UCSF Gladstone Genomic Core and the UCSF Center for Advanced Technology). The data were analyzed using 10x Genomics Cell Ranger 6.0.1 with a recovery of 4000-7000 cells per experiment (see Supplementary Table 1).

### Chromatin Immunoprecipitation sequencing studies

#### Fixed

The following antibodies were used for crosslinked chromatin immunoprecipitation studies: CTCF antibody pool (equal ng per antibody: Santa Cruz sc-271514, sc-271474, Bethyl Laboratory A300-542A, Millipore 07-729), Rad21 antibody pool (equal ng per antibody: Abcam ab992, Thermo Fisher PA528344, Bethyl Laboratory A300-080A). With the exception of ChIP-Seq experiments for CTCF performed in OSNs where about 1 million sorted cells were used per IP, about 2 to 5 million cells were used. Cells were crosslinked with 1% formaldehyde for 10 minutes at room temperature. Formaldehyde was quenched by adding glycine to a final concentration of 0.125 M for 5 minutes at room temperature. Cells were then washed twice with 1X cold PBS with protease inhibitors twice and pelleted. Cell pellets were stored at -80°C till use. Cells were lysed in lysis buffer (50 mM Tris pH 7.5, 140 mM NaCl, 0.1% SDS, 0.1% sodium deoxycholate, 1% Triton X-100) for 10 minutes. Nuclei were spun for 10 minutes at 1000 x g and resuspended in the sonication buffer (10 mM Tris pH 7.5, 0.5% SDS) as 5 million nuclei per 300 μl sonication buffer. Chromatin was sheared by Covaris (Peak power 105.0; Duty Factor 2.0; Cycle/Burst 200; Treatment 960 sec; Temperature 4-8°C). Following a spin at 13,000 x g for 10 minutes to remove debris, the sheared chromatin was diluted such that the final binding buffer concentration was 15 mM Tris-HCl pH 7.5, 150 mM NaCl, 1 mM EDTA, 1% Triton X-100, 0.1% SDS) and incubated for 2 hours with dynabeads G pre-equilibrated in the binding buffer for pre-clearing of the chromatin. Post-cleared chromatin was then incubated with the specific antibody overnight (1 μg of antibody was used per 5 million nuclei). The next day, dynabeads G were added to the chromatin-antibody mix for 2 hours. A total of four washes with 1X wash buffer (100 mM Tris pH 7.5, 500 mM LiCl, 1% NP-40, 1% sodium deoxycholate) and one wash with TE buffer (10 mM Tris pH 7.5, 1 mM EDTA) were performed. The elution was performed at 65°C for 1 hour in the elution buffer (1% SDS, 250 mM NaCl). All steps, with the exception of the elution, were performed at 4°C. All buffers, with the exception of the TE and elution buffer, contained 1X protease inhibitors. The eluted chromatin was reverse-crosslinked overnight at 65°C and the DNA was purified with AMPure XP beads (Beckman Coulter).

#### Native

The following antibodies were used for native chromatin immunoprecipitation studies: Histone H3 Lysine 9 tri-methyl (Abcam ab8898). Histone H3 Lysine 4 tri-methyl (ThermoFisher PA5-27029), Histone H3 Lysine 27 acetylation (Abcam ab4729). About 100,000 to 200,000 sorted OSNs were lysed and the isolated nuclei were subjected to MNase treatment. The MNase-treated chromatin was then incubated with the corresponding antibody overnight. The next day, dynabeads G and A were added to the chromatin-antibody mix for 2 hours. A total of four washes with 1X wash buffer I (50 mM Tris pH 7.5, 125 mM NaCl, 0.1% Tween 20, 10 mM EDTA) and three wash with 1X wash buffer II TE buffer (5 mM Tris pH 7.5, 17.5 mM NaCl, 0.01% NP-40, 1 mM EDTA) were performed. The elution was performed at RT for 1 hour in the elution buffer (1% SDS, 250 mM NaCl, 2 mM DTT). All steps, with the exception of the TE buffer and elution buffer, were performed at 4°C. All buffers, with the exception of the elution buffer, contained 1X protease inhibitors. For IPs involving H3K27ac, sodium butyrate (Na-But) was supplemented to all the buffers.

Libraries for both fixed and native ChIP-Seq were prepared using the Ovation Ultralow V2 DNA-Seq Library Preparation Kit (TECAN). The quality of the libraries was assessed by bioanalyzer and quantified using a combination of bioanalyzer and qubit. Libraries were sequenced on a NEXT-Seq 500/550 (UCSF Center for Advanced Technology).

### *In situ* Chromatin Capture Conformation (Hi-C)

About 500,000 sorted OSNs were lysed and intact nuclei were processed through an *in situ* Hi-C protocol as previously described with a few modifications ^37^. Briefly, cells were lysed with lysis buffer (50 mM Tris pH 7.5 0.5% Igepal, 0.25% Sodium-deoxycholate, 0.1% SDS, 150 mM NaCl, and protease inhibitors). Pelleted intact nuclei were then resuspended in 0.5% SDS and incubated for 20 minutes at 65°C for nuclear permeabilization. After quenching with 1.1% Triton-X for 10 minutes at 37°C, nuclei were digested with 6 U/μl of DpnII in 1x DpnII buffer overnight at 37°C. Following initial digestion, a second DpnII digestion was performed at 37°C for 2 hours. DpnII was heat-inactivated at 65°C for 20 minutes. For the 1.5 hours fill-in at 37°C, biotinylated dATP (Jena Bioscience) was used instead of dATP to increase ligation efficiency. Ligation was performed at 25°C for 4 hours. Nuclei were then pelleted and sonicated in sonication buffer (10 mM Tris pH 7.5, 1 mM EDTA, 0.25% SDS) on a Covaris S220 (Peak power 105.0; Duty Factor 2.0; Cycle/Burst 200; Treatment 960 sec; Temperature 4-8°C). DNA was reverse cross-linked overnight at 65°C with proteinase K and RNAse A. Reverse cross-linked DNA was purified with 2x AMPure beads following the standard protocol. Biotinylated fragments were enriched using Dynabeads MyOne Streptavidin T1 beads. The biotinylated DNA fragments were prepared for next-generation sequencing on the beads by using the Nugen Ovation Ultralow kit protocol with some modifications. Following end repair, magnetic beads were washed twice at 55°C with 0.05% Tween, 1 M NaCl in Tris/EDTA pH 7.5. Residual detergent was removed by washing the beads twice in 10 mM Tris pH 7.5. End repair buffers were replenished to original concentrations, but the enzyme and enhancer was omitted before adapter ligation. Following adaptor ligation, beads underwent five washes with 0.05% Tween, 1 M NaCl in Tris/EDTA pH 7.5 at 55°C and two washes with 10mM Tris Ph 7.5. DNA was amplified by 10 cycles of PCR, irrespective of starting material. Beads were reclaimed and amplified DNA fragments were purified with 0.8x AMPure beads. Quality and concentration of libraries were assessed by Agilent Bioanalyzer and Qubit. Samples were sequenced paired-end on a NEXT-Seq 500/550 or NovaSeq (UCSF Gladstone Genomic Core and the UCSF Center for Advanced Technology).

### Whole Genome Bisulfite Sequencing

Bisulfite DNA reactions and library preparations were performed using the Ultralow Methyl-Seq with TrueMethyl oxBS module, Nugen, following the steps indicated by the protocol. Genomic DNA (gDNA) was isolated from 100,000 sorted OSNs using the PureLink Genomic DNA Kit. A total of 100ng of purified gDNA was used for each experiment. Two independent biological experiments were performed per genotype. The quality of the libraries was assessed by bioanalyzer and quantified using a combination of bioanalyzer and qubit. Libraries were sequenced on a Nova-Seq (UCSF Center for Advanced Technology).

### Bioinformatic Analysis of Sequencing Data

For RNA-Seq experiments, raw FASTQ files were aligned with STAR using the mm10 reference genome. The initial 4 base pairs of both paired reads were trimmed prior to alignment.

For ChIP-Seq experiments, raw FASTQ files were aligned using Bowtie2 using the mm10 reference genome upon adapter sequences removal using CutAdapt. Reads from the 3xCBS FACS-sorted OSNs were aligned to a modified mm10 including the inserted sequence. For fixed ChIP-Seq, only uniquely aligning reads were selected and further processed.

For *in situ* Hi-C experiments, raw FASTQ files were processed using HiC-Pro (see Supplementary Table 2) ^38^. Hi-C pyramid plots and difference maps were generated using GENOVA ^39^.

For WGBS experiments, analysis was performed by following the steps indicated here: https://github.com/nugentechnologies/NuMetWG.

### Preparation of Mouse Brain Tissue Sections

Animals were euthanized by CO_2_ induction followed by cervical dislocation. Brains were immediately dissected after euthanasia. For experiments using olfactory bulb (OB) tissue, the OB was dissected and immersion fixed in 4% paraformaldehyde (PFA) for 8 min, then washed in PBS 3×5 min at RT and incubated in 30% sucrose overnight at 4°C. The following day, samples were incubated in a 1:1 30% sucrose and OCT solution for 1 hour, then moved to OCT for 1 hour prior to freezing. Samples were frozen in embedding molds using isopropanol and dry ice. For experiments targeting 5-HT neurons in the raphe and in the hippocampus, whole brains were dissected and placed in 4% PFA overnight at 4°C. The brains were then washed in PBS 3×5 min at RT and transferred to 15% sucrose until they sunk, and subsequently to 30% sucrose at 4°C overnight. Once sunk, OB tissue was removed from 30% sucrose and placed in a tube containing 1:1 solution of 30% sucrose and OCT embedding compound (Sakura, Ref 4583) for 1 hour incubation at RT, followed by another hour of RT incubation in an embedding mold containing pure OCT. Whole brains were directly transferred from 30% sucrose solution to an embedding mold containing 2:1 30% sucrose and OCT. To freeze the tissue in its embedding solution, the mold was flash frozen in dry ice and isopropanol. Tissue was stored at -80°C until sectioning.

Samples in OCT were placed in a Leica cryostat (Leica Microsystems, Wetzlar, Germany) 30-45 mins before sectioning to equilibrate. OCT was used to freeze samples to cryostat chucks. Sectioning was performed with a chamber temperature of -21°C and an object temp of -17°C. Samples were sectioned coronally at 16 μm (OB) and at 20 μm (hippocampus and raphe). All sections were mounted onto superfrost plus slides (Fisher Scientific, Cat. No. 12-550-15) and stored at -20°C until downstream use.

### Immunohistochemistry

For immunostaining of the OB tissue, sections were fixed with 4% PFA in PBS, washed 3x 5 minutes with PBS containing 0.1% Triton-X100 and blocked in 4% donkey serum in PBS + 0.1% Triton-X100 for 30 minutes at room temperature. For immunostaining of the hippocampus region, sections were directly washed with PBS + 0.1 Triton-X100 and incubated with the same blocking serum for 30 minutes at room temperature. Following the blocking step, the slides were incubated in a humid chamber with primary antibodies diluted in blocking serum for 24 hours at 4°C. We used the following primary antibodies: rabbit anti-SERT (1:500, Millipore Sigma, PC177L100UL) and guinea pig anti-Vglut2 (1:2000, Millipore Sigma, AB2251-I). After extensive washing with PBS + 0.1% Triton-X100, sections were incubated with Alexa dye–conjugated secondary antibodies diluted in blocking serum for 2 hours before being washed again and mounted with Vectashield Antifade Mounting Medium with DAPI (Vector Laboratories). Alexa Fluor® 488 Donkey IgG, anti-rabbit (1:2000, Invitrogen, A21206) and Alexa Fluor 647 Goat anti-Guinea Pig IgG (1:2000, Invitrogen, A21450) were used as secondary antibodies.

### Hybridization Chain Reaction Experiments on Fixed Tissue Sections

RNA FISH experiments were performed using hybridization chain reaction (HCR) technology from Molecular Instruments. Fixed tissue sections were taken from -20°C and allowed to dry at RT for 5 min. They were then washed with PBS for 5 min to remove excess OCT. An HCR protocol (dx.doi.org/10.17504/protocols.io.b2frqbm6) was then performed using the following probes and matching hairpins ordered from Molecular Instruments: Tryptophan hydroxylase 2 (Tph2), lot number PRE854 and Protocadherin alpha-C2 (Pcdhαc2), lot number PRK987. The hairpin amplification systems used for Tph2 and Pcdhαc2 were B1 at 546 nm and B3 at 647 nm, respectively.

### Imaging studies

Sections from the OB and raphe were imaged using a Yokogawa CSU-W1 SoRa spinning disk confocal. To improve Pcdhαc2 mRNA visualization, images of the raphe were deconvoluted using the Huygens Essential software application. Z stacks of 11 images with steps of 0.25 μm were used for the deconvolution, and maximum intensity projections of the resulting deconvolution images were chosen for visualization. No deconvolution or z stack projection was used for images subjected to puncta quantification (see “Quantification of Pcdhαc2 Expression”). Sections from the hippocampus were imaged using a Leica DMi8 inverted microscope and stitched using the LAS X software platform. All images were post processed in FIJI.

### Quantification of Pcdhαc2 Expression

Pcdhαc2 expression in individual 5-HT neurons was quantified using a custom CellProfiler pipeline ^40^. The pipeline categorized input images from HCR experiments into groups of three as follows: 1) Pcdhαc2 mRNA signal in the 647 nm channel; 2) Tph2 mRNA signal in the 561 nm channel; and 3) manual segmentations of 5-HT somas based on Tph2 expression performed in FIJI. The pipeline first identified 5-HT somas from the manual segmentations using the two-classes Otsu thresholding method with a correction factor of 1.0. These somas were then used as a mask over the Pcdhαc2 puncta in the 647 nm image, only keeping puncta residing within 5-HT somas. The masked Pcdhαc2 puncta image was then enhanced for speckle features with a maximum size of 10 pixels to make mRNA puncta easier to identify. Pcdhαc2 puncta were identified using the Robust Background thresholding method with a correction factor of 1.0, lower and upper outlier fractions of 0.05, and two standard deviations. Clumped puncta were distinguished by shape and dividing lines between clumped objects were drawn based on intensity. Finally, the RelateObjects module was used to assign each Pcdhαc2 punctum to its parent 5-HT soma. Data was exported for the number of Pcdhαc2 puncta within each 5-HT soma, 5-HT soma area, 5-HT soma Tph2 signal intensity, and Pcdhαc2 puncta intensity.

### Quantification of Glomerular Structures

Distinct glomeruli were defined as spherical Vglut2+ structures bordered by periglomerular cell nuclei. Number of distinct glomeruli were quantified from 6 evenly spaced coronal sections along the antero-posterior axis of the olfactory bulb. To minimize bias introduced by tissue damage, only medial glomeruli were included in the quantifications, but differences in glomerular structure were reported across the entire antero-posterior axis of the olfactory bulb.

## QUANTIFICATION AND STATISTICS

The statistical tests in this study were performed by Student’s unpaired *t*-test to determine statistical significant effects (p < 0.05).

## ACKNOWLEDGMENTS

We thank Dr. Tom Maniatis, Dr. Stavros Lomvardas, Dr. Geeta Narlikar, Dr. Elphege Nora, Dr. Hiten Madhani, Dr. Vijay Ramani, Dr. Mercedes Paredes, Dr. George Mountoufaris, Dr. Enrico Cannavo’, Hani Shayya and members of the Canzio lab for helpful discussions and suggestions. We thank Chiamaka Nwakeze for the initial training of S.R. in the generation of transgenic animals. We thank Dr. Luca Giorgetti for providing us with the DNA sequence utilized to engineer the 3xCBS mouse. We thank the UCSF Gladstone Mouse Transgenic Core for help in the generation of transgenic animals and Mylinh Bernardi at the UCSF Gladstone Genomic Core and the UCSF Center for Advanced Technology for help in sequencing experiments. We thank SoYeon Kim and the UCSF Center for Advanced Light Microscopy for support with image acquisition and analysis. This work was funded by the National Institute of General Medicine, grant R00 GM121815-05 (D.C.), the National Institute of Mental Health, grant DP2 MH129955-01 (D.C.) and the National Science Foundation, Graduate Research Fellowship Grant No. 2034836 (A.B.).

## AUTHOR CONTRIBUTIONS

D.C. conceived the project and assisted with the design and interpretation of the experiments. L.K. designed, performed, and interpreted the single cell RNA-Seq experiments, analyzed all the Hi-C datasets, and helped with the conceptualization of the genomic data within the story. G.F.S. performed the bulk of the genomic experiments. S.M.R. generated all the unpublished mouse lines and performed their initial characterization with the help of E.S.C.. J.L. designed, performed, and interpreted the immunohistochemistry studies in the olfactory bulb. A.C. designed, performed, and interpreted the immunohistochemistry studies in the hippocampus. A.B. designed, performed, and interpreted the RNA FISH studies. A.H. trained D.C. on performing Hi-C experiments. M.H.M. assisted in mouse colony management. D.C. wrote the paper with the substantial help from all authors.

## COMPETENTING INTERESTS

Authors declare no competing interests.

## MATERIALS and CORRESPONDENCE

All data are available in the main text and in the supplementary materials and can be made available from the corresponding author upon request. Source data have been deposited in NCBI’s Gene Expression Omnibu and are accessible through GEO Series accession number GSE191195.

## TABLES

**Table 1:** Single-cell RNA-seq analysis

| Sample Name | Cell type | Total Raw Reads | Mapping rate to mm10 | Est. # of cells from Cellranger | Total genes detected | Median Genes per cell | Median UMI count per cell |
| --- | --- | --- | --- | --- | --- | --- | --- |
| S1 | OSN WT | 293179627 | 95.10% | 6476 | 20476 | 3928 | 11122 |
| S2 | OSN 3xCBS | 274530522 | 95.00% | 6393 | 20327 | 3960 | 10622 |
| S3 | OSN HS5-1i | 193216267 | 95.00% | 4150 | 19224 | 3972 | 11223 |
| S4 | OSN Rad21cKO | 196840615 | 94.50% | 5213 | 21858 | 3816 | 10636 |

**Table 2:**
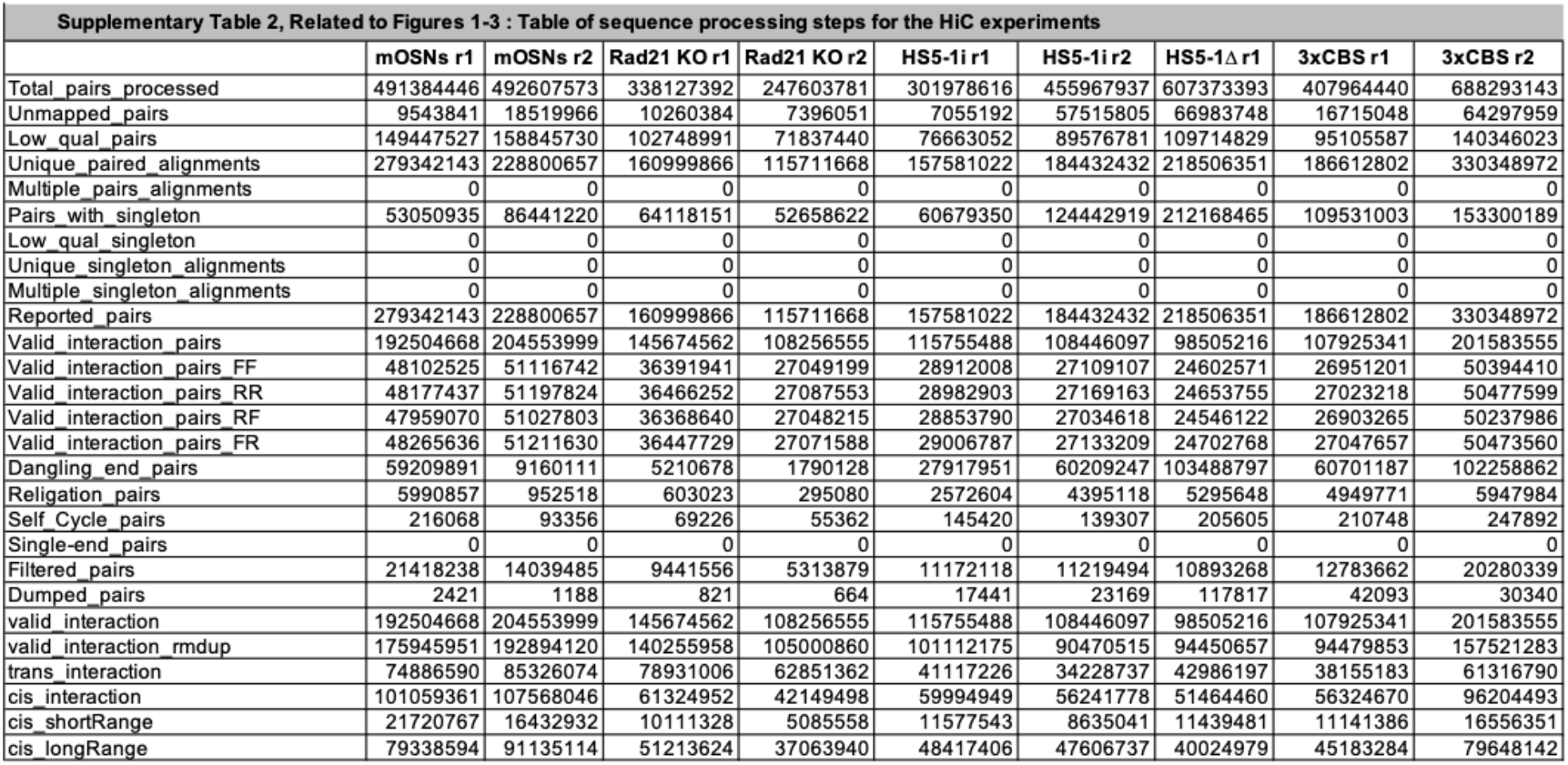
HiC analysis

**Table 3:**
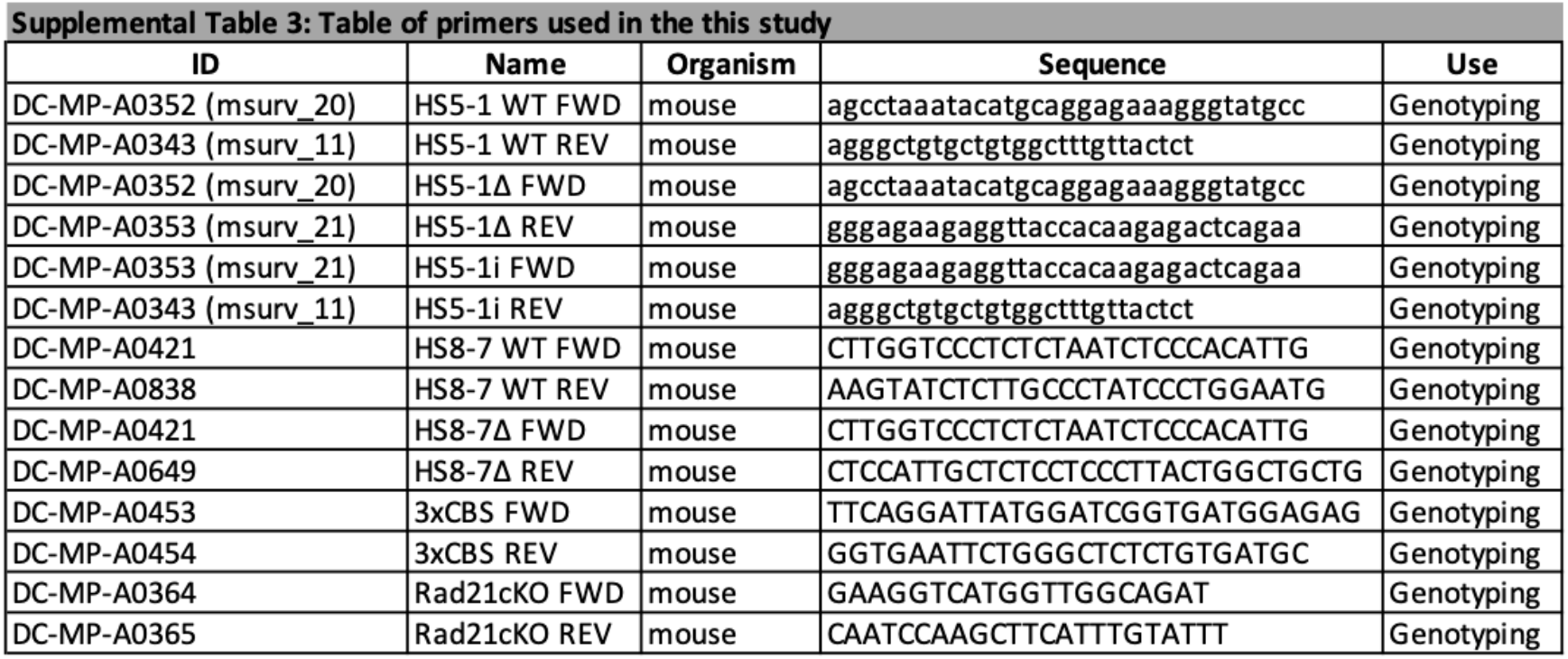
Primer sequences

**Extended Data Figure 1.**
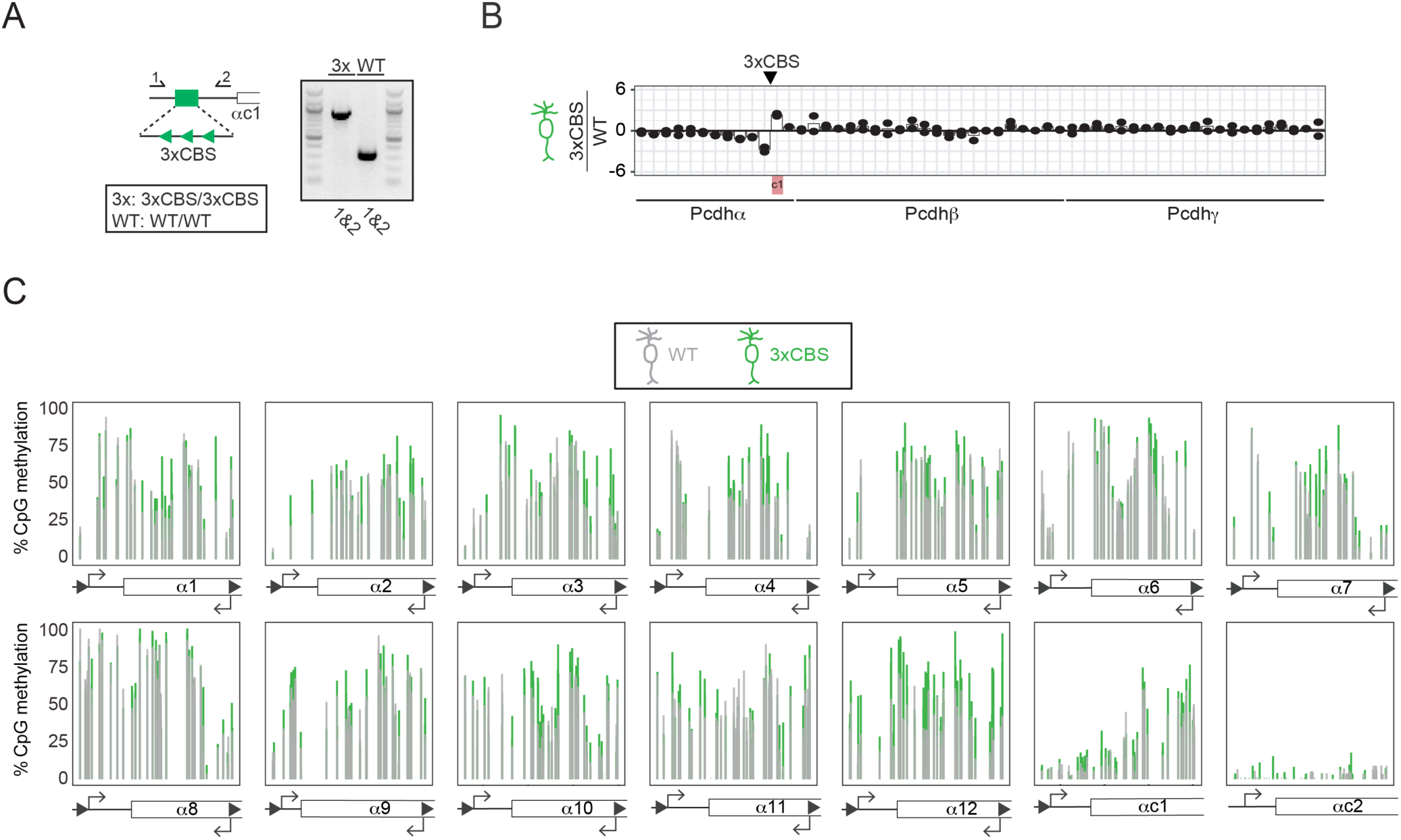
Generation of the 3xCBS mouse and its effect on Pcdh gene expression regulation. **(a)** Schematic of the engineered 3xCBS mouse and its genotyping (Primer numbers 1=A0453, 2=A0454, see Supplemental Table 3). **(b)** Relative RNA-seq changes upon ectopic 3xCBS insertion compared to WT OSNs of all clustered Pcdh isoforms. Arrowhead indicates the location of the 3xCBS. **(c)** CpG methylation across the Pcdhα promoters in WT (grey) and 3xCBS (green) OSNs. For b and c, two biological replicates are shown.

**Extended Data Figure 2.**
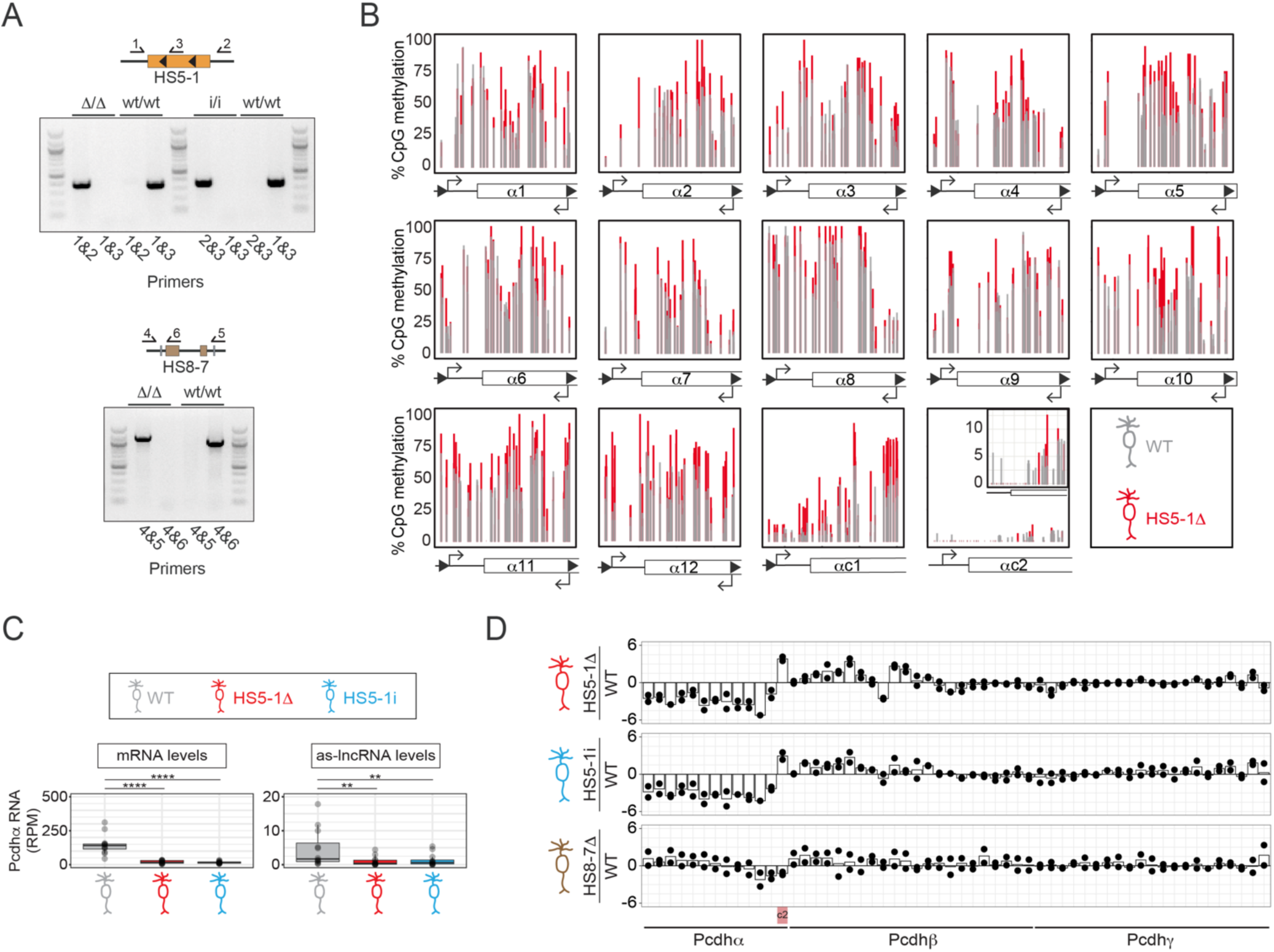
Cohesin extrusion regulates the activity of the HS5-1 enhancer. **(a)** Schematics of the engineered HS5-1Δ, HS5-1i and HS8-7Δ mice and their genotyping (primer numbers 1=A0352, 2=A0353, 3=A0343, right, 4=A0421, 5=A0838, 6=A0649, see Supplemental Table 3). **(b)** CpG methylation across the Pcdhα promoters WT (grey) and HS5-1Δ (red) OSNs. **(c)** Quantification of sense and antisense RNA levels of Pcdhα isoforms in HS5-1Δ and HS5-1i OSNs compared to WT. **(d)** Relative RNA-seq changes upon HS5-1Δ, HS5-1i and HS8-7Δ compared to WT OSNs of all clustered Pcdh isoforms. For b, c and d, two biological replicates are shown.

**Extended Data Figure 3.**
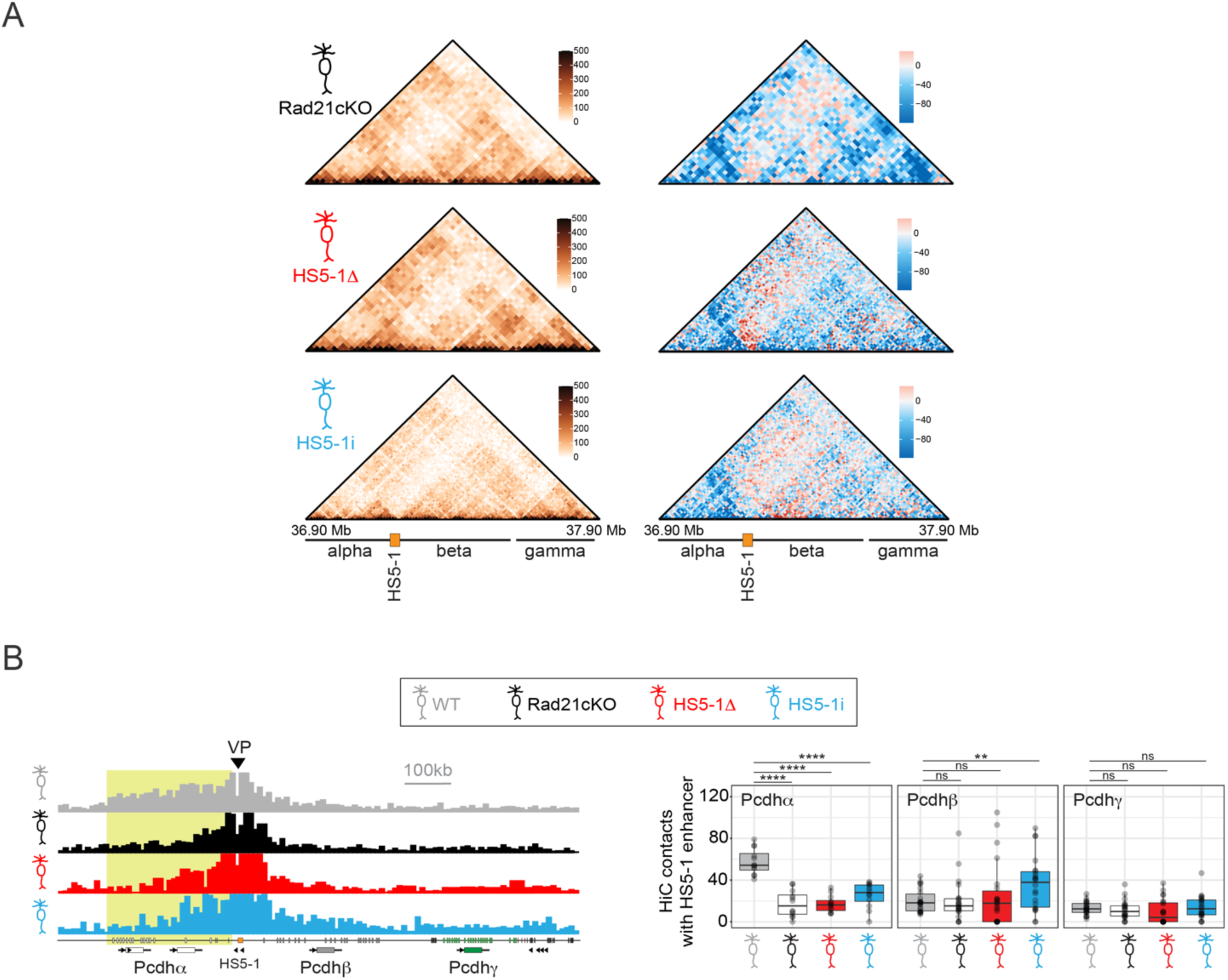
Chromatin architecture of the Pcdh locus upon disruption of DNA loop extrusion. **(a)** (Left) HiC maps of the Pcdh locus from Rad21cKO, HS5-1Δ and HS5-1i OSNs. (Right) Difference map with gained interactions compared to WT in red, lost interactions in blue. **(b)** Left: Virtual 4C of the chromatin contacts from the HS5-1. Viewpoint indicated by arrowheads. Right: Quantification of HiC chromatin interactions between Pcdh α, β and γ promoters and the HS5-1 enhancer in Rad21cKO, HS5-1Δ, HS5-1i OSNs compared to WT.

**Extended Data Figure 4.**
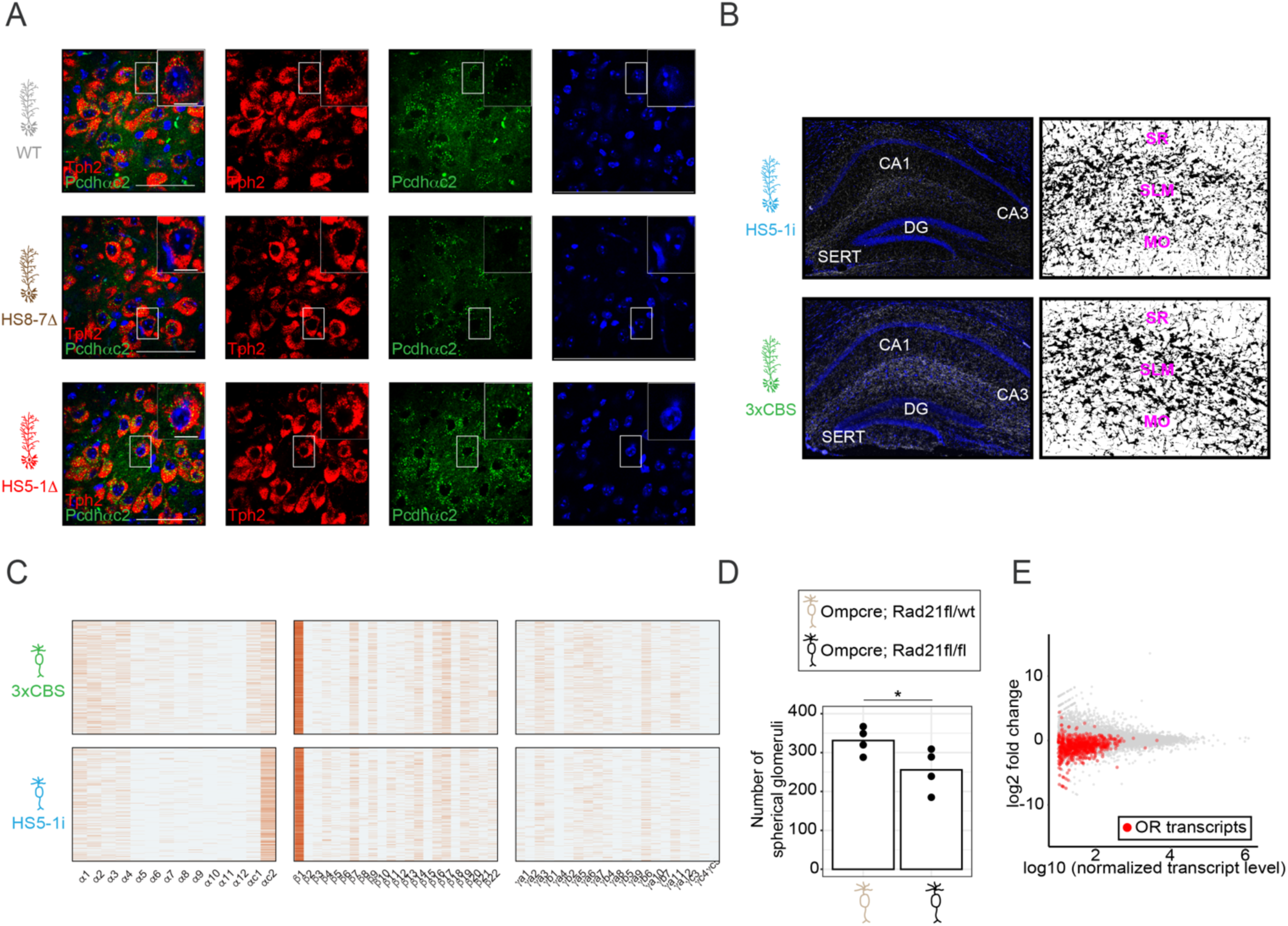
Cohesin-dependent and -independent mechanisms of Pcdh promoter choice in 5-HTs and OSNs. **(a)** RNA FISH in the raphe isolated from WT, HS8-7Δ and HS5-1Δ mice. Tph2, red and Pcdhαc2, green. DAPI in blue. Scale bars: 50 μm. **(b)** 5-HT wiring in HS5-1i and 3xCBS mice. Left: 5-HTs are indicated by IHC against SERT. SERT, white; DAPI, blue. CA1, CA3 and DG refer to cornu ammonis 1 and 3 and dentate gyrus, respectively. Right: Binary mask of SERT signal in presented images. SR, SLM and MO refer to stratum radiatum, stratum lacunosum moleculare, and molecular layer. **(c)** Single-cell Pcdh promoter choice from all three clusters from individual OSNs isolated from 3xCBS and HS5-1i OSNs. Each row indicates a single cell, each column a single isoform. **(d)** Quantification of the number of spherical glomeruli from six evenly distributed sections across the antero-posterior axis of the olfactory bulb (n=4 for each genotype). **(e)** RNA-seq analysis of transcript levels in omp-cre;Rad21fl/fl (Rad21cKO) compared to omp-cre;Rad21fl/wt (Rad21het). OR transcripts are shown in red.

